# Antagonism of GluK1-containing Kainate Receptors Reduces Ethanol Consumption by Modulating Ethanol Reward and Withdrawal

**DOI:** 10.1101/2021.04.24.441015

**Authors:** Natalia A. Quijano Cardé, Erika E. Perez, Richard Feinn, Henry R. Kranzler, Mariella De Biasi

**Author notes:** Department of Psychology, Xavier University of Louisiana, New Orleans, LA 70125. Corresponding author: Mariella De Biasi.

## Abstract

Alcohol use disorder (AUD) is a neuropsychiatric condition affecting millions of people worldwide. Topiramate (TPM) is an antiepileptic drug that has been shown to reduce ethanol drinking in humans. However, TPM is associated with a variety of adverse effects due to its interaction with many receptor systems and intracellular pathways. Thus, a better understanding of the role of TPM’s main molecular targets in AUD could yield better therapeutic tools. GluK1-containing kainate receptors (GluK1*KARs) are non-selectively inhibited by TPM, and genetic association studies suggest that this receptor system could be targeted to reduce drinking in AUD patients. We examined the efficacy of LY466195, a selective inhibitor of GluK1*KAR, in reducing ethanol consumption in the intermittent two-bottle choice paradigm in mice. The effect of LY466195 on various ethanol-related phenotypes was investigated by quantification of alcohol intake, physical signs of withdrawal, conditioned place preference (CPP) and *in vivo* microdialysis in the nucleus accumbens. Selective GluK1*KAR inhibition reduced ethanol intake and preference in a dose-dependent manner. LY466195 treatment attenuated the physical manifestations of ethanol withdrawal and influenced the rewarding properties of ethanol. Interestingly, LY466195 injection also normalized changes in dopamine levels in response to acute ethanol in ethanol-dependent mice, but had no effect in ethanol-naïve mice, suggesting ethanol state-dependent effects. The data point to GluK1*KARs as an attractive pharmacological target for the treatment of AUD.

## 1. INTRODUCTION

Alcohol use disorder (AUD) is a highly prevalent, heterogeneous neuropsychiatric disorder that results from the interplay of social, environmental, and genetic factors. More than 17 million Americans meet criteria for current AUD, but less than 20% of those individuals ever receive treatment (Cohen et al., 2007; Grant et al., 2015). Currently approved medications for AUD (Akbar et al., 2018) have modest effects on drinking outcomes and are not widely prescribed, underscoring the need for more efficacious medications for treating AUD.

Antipsychotics, antidepressants, and anticonvulsants have also shown potential for treating AUD (Soyka and Müller, 2017). The anticonvulsant topiramate (TPM) was shown in randomized, placebo-controlled trials to significantly reduce drinking in alcohol-dependent patients (Blodgett et al., 2014), and the same effect was observed in rodent studies (Breslin et al., 2010; Nguyen et al., 2007; Zalewska-Kaszubska et al., 2013). Interestingly, the effects of TPM seem to be specific to the ethanol-dependent state, as the drug does not affect the reinforcing or behavioral effects of acute ethanol (Chen and Holmes, 2009; Gremel et al., 2006). Additional studies suggest that TPM reduces ethanol consumption in part by reducing animals’ motivation for alcohol in a progressive-ratio model of self-administration (Hargreaves and McGregor, 2007). TPM also reduces anxiety-like behavior during ethanol withdrawal (Farook et al., 2007). Thus, TPM may reduce drinking by affecting both the positive and negative reinforcing properties of ethanol.

TPM has several pharmacological actions, including antagonism of the kainate-subtype of glutamate receptors (KARs) (Gibbs et al., 2000; Williams et al., 2000). TPM displays higher selectivity and potency for KARs containing GluK1 and GluK2 subunits, encoded by *GRIK1* and *GRIK2*, respectively (Gryder and Rogawski, 2003; Kaminski et al., 2004). The potential targeting of GluK1-containing (GluK1*) KARs by TPM is of particular interest, because the major (C) allele of the single nucleotide polymorphism (SNP) rs2832407, a C-to-A non-coding intronic substitution in *GRIK1*, was associated with alcohol dependence (Kranzler et al., 2009). Moreover, a neuroimaging study showed that the rs2832407*C was associated with neuronal activation in brain regions positively correlated with subjective alcohol craving and relapse risk in abstinent, alcohol-dependent individuals (Bach et al., 2015). The SNP also moderated responses to TPM in treating AUD, with reduced drinking seen only among *GRIK1*\*rs2832407 C-allele homozygotes (rs2832407*CC) (Kranzler et al., 2014a). In that study, rs2832407*CC individuals showed persistent reductions in heavy drinking during the six months following the cessation of pharmacotherapy (Kranzler et al., 2014b). In a study of heavy drinkers, the rs2832407*CC genotype was also associated with less severe TPM-induced adverse effects (Ray et al., 2009). Although the functional significance of this *GRIK1* SNP is unclear, the findings suggest that GluK1*KARs play an important role in AUD risk and treatment. TPM causes a variety of mild-to-moderate adverse effects (Johnson et al., 2007; Knapp et al., 2015) necessitating a slow dose titration treatment initiation which limits its utility in treating AUD. Thus, selective inhibition of GluK1*KARs could be efficacious in decreasing drinking with fewer adverse effects than TPM.

The GluK1 subunit is expressed throughout the brain, including in regions that modulate different aspects of addiction, such as anterior cingulate cortex (Wu et al., 2007), hippocampus (Christensen et al., 2004; Clarke and Collingridge, 2004), ventral pallidum (Bischoff et al., 1997; Wisden and Seeburg, 1993), basolateral amygdala (BLA) (Wu et al., 2007), and cortical afferents projecting to the nucleus accumbens (NAc) (Bettler et al., 1990; Casassus and Mulle, 2002). Hence, GluK1*KARs could modulate various aspects of alcohol dependence, such as positive reinforcement, withdrawal, and alcohol-seeking behaviors. Acute ethanol inhibits GluK1*KAR-mediated neurotransmission in the BLA (Läck et al., 2008), whereas chronic ethanol exposure promotes a hyperglutamatergic state and increases GluK1*KAR activation (Läck et al., 2009). Chronic, ethanol-mediated GluK1*KAR-modulation of BLA glutamatergic function also suggests that these receptors might contribute to the aversive components of ethanol withdrawal, especially given the crucial role of this brain region in the pathophysiology of anxiety in the context of addiction (Sharp, 2017).

Based on these findings, we sought to determine whether selective GluK1*KAR inhibition reduces ethanol intake and preference. In addition, we examined the role of GluK1*KARs in ethanol reward and withdrawal. Lastly, we measured the effect of GluK1*KAR antagonism on ethanol-induced changes in dopamine (DA) levels in the NAc during various ethanol-associated states. Our study examined the effects of LY466195, a decahydroisoquinoline that acts as competitive GluK1*KAR antagonist (Ki = 52 ± 22 nM) with a greater than 100-fold selectivity for GluK1*KARs versus the other kainate and AMPA receptor subtypes (Weiss et al., 2006).

## 2. MATERIALS AND METHODS

### 2.1. Animals

We examined 2- to 6- month-old C57Bl/6J mice from a colony maintained in the laboratory. Male and female mice were included in all the experiments. On average, across all experiments and experimental groups, males represented 50.6±15.0% (mean standard ± deviation) of the experimental subjects. Throughout the study, mice were housed in standard ‘shoebox’ cages (7.6 in × 5.1 in × 15 in) with 1 cm corn cob bedding, a cotton nestlet, and *ad libitum* access to water and food (Labdiet 5053, PMI, Brentwood, MO). The mice were housed in a temperature- and humidity-controlled room (65-75 °F, 40-60% relative humidity) with a 12-h light/dark cycle. For consumption studies, experimental subjects were single-housed with a tent-shaped cardboard house (Shepherd shack) for additional environmental enrichment, and the experiments began after one week of acclimation to the housing conditions.

### 2.2. Ethical Statement

All procedures were approved by the University of Pennsylvania Institutional Animal Care and Use Committee. This study followed the guidelines of the NIH Guide for the Care and Use of Laboratory Animals.

### 2.3. Materials

Ethanol solutions were prepared by diluting 190 proof ethanol (Decon Labs, Inc., King of Prussia, PA) in filtered water (for drinking experiments) or phosphate buffered saline (PBS, for injections). The selective GluK1*KAR antagonist LY466195 was provided by Eli Lilly & Co., Indianapolis, IN, USA (Weiss et al., 2006). LY466195 solutions were prepared by dissolving LY466195 in PBS. Chemicals used for *in vivo* microdialysis were from Sigma-Aldrich (Millipore Sigma, St. Louis, MO).

### 2.4. Intermittent Two-bottle Choice Drinking Paradigm

The intermittent-two-bottle choice (I2BC) drinking paradigm promotes high voluntary ethanol intake in mice (Hwa et al., 2011; Melendez, 2011; Rosenwasser et al., 2013).

Briefly, fluids were administered using 50-mL bottles with straight stainless-steel, open-tip tubes. Following a one-week habituation period, mice began to receive ethanol on Mondays, Wednesdays and Fridays for 24 h, starting 3 hours after lights off. During periods of ethanol deprivation, they had *ad libitum* access to filtered water in both bottles. During the first week of alcohol exposure, mice were provided with increasing ethanol concentrations (3, 6, and 10% v/v) in one of the bottles, followed by 20% (v/v) ethanol for the rest of the experiment. Mice were maintained in the I2BC for at least six weeks before LY466195 treatment, and the position of the bottles was alternated to avoid side preference. Bottles were weighed to the nearest hundredth to determine 2-h and 24-h ethanol and water consumption. Empty cages containing ethanol and filtered water bottles were used to determine changes in bottle weight due to evaporation and leakage due to cage movement. Mice were weighed at least once weekly to determine ethanol dose (g/kg). Ethanol consumption was calculated using ρ_solution_=1 g/mL and ρ_ethanol_=0.789 g/mL. Ethanol preference was calculated as the ratio of ethanol to total fluid intake (mL ethanol solution/mL total fluid).

In our hands, blood plasma ethanol concentrations reach values of 15.6 ± 4.3 mg/dL and 16.0 ± 4.8 mg/dL in C57BL/6J males and females mice, respectively, when mice are allowed to drink 20% ethanol for 2 hrs in the I2BC paradigm.

### 2.5. Effect of LY466195 on Ethanol Consumption

Mice were habituated to receive intraperitoneal (i.p.) saline injections (0.01 mL/g) before I2BC ethanol presentation for a minimum of two weeks.

To study short-term withdrawal, mice undergoing a 24-h period of ethanol withdrawal received an i.p. injection of LY466195 (0, 4, 10, 20, or 40 mg/kg) immediately before ethanol presentation. The injection volume was maintained at 0.01 mL/g for all doses. Mice were treated with multiple LY466195 doses in a randomized order with at least one drinking session between each LY466195 dose.

For protracted withdrawal, a subset of mice from the short-term withdrawal experiment was exposed to ethanol only (no LY466195 administrations) in the I2BC for 2.5 weeks before they were examined in the paradigm described by Melendez et al., 2006 with modifications. Briefly, mice with stable ethanol consumption underwent three 6-day cycles of prolonged withdrawal. Each Friday, mice received an i.p. injection of 20 mg/kg LY466195 or saline, and ethanol was immediately reintroduced via the two-bottle choice paradigm for 24 h, as previously described. Throughout the three cycles, mice were maintained on the same drug treatment. Ethanol dose and preference were determined as described for the I2BC paradigm.

### 2.6. Physical Signs of Withdrawal

We compared ethanol-naïve mice to mice with stable consumption in the I2BC with respect to signs of withdrawal. Ethanol-dependent mice were tested 24 h after the last ethanol exposure (short-term withdrawal) or 4 h into the drinking session (satiety) for physical signs of withdrawal [e.g., shaking, scratching, grooming, chewing, paw tremors, and cage scratching (Perez et al., 2015; Perez and De Biasi, 2015)]. To examine physical behaviors, mice were tested in standard housing cages with 1 cm corn cob bedding during the dark phase of the light cycle in a behavioral room with sound-attenuating walls and dim lights (~3 lux). Subjects were habituated to the testing room for 30 min before behavioral testing. After the habituation period, physical signs were monitored for 20 min, followed by an i.p. injection of either saline or 20 mg/kg LY466195. Fifteen min after LY466195 administration, physical signs were scored again for 20 min. The physical behaviors were scored with 360-degree visibility of the experimental cage. Auditory inputs are needed for the quantification of certain physical behaviors (i.e. vocalizations, cage scratching, chewing), thus all behaviors were scored in real-time. Individual physical signs and total scores (i.e. summation of the occurrence of each sign category per mouse) were examined. The percentage change was calculated as follows: [(total_post-test_ – total_pre-test_)/(total_pre-test_)]×100%.

### 2.7. Conditioned Place Preference (CPP)

We administered LY466195 to ethanol-naïve mice and mice with stable consumption in the I2BC paradigm to test its effect on the acquisition of ethanol CPP (Gremel et al., 2006; Nocjar et al., 1999). The CPP apparatus consisted of two equal-sized chambers (15 cm × 22 cm each) with distinct visual and tactile cues. For the visual cues, each chamber contained different black-and-white patterns on the walls (i.e. stripes vs circles). For the tactile cues, the chambers were lined with removable floors made of clear, ribbed shelf liner (“striped”) or white plastic canvas (“grid”). The testing apparatus and the removable floors were sanitized with 70% ethanol before each testing and conditioning session. Mice were habituated to handling and saline i.p. injections for a week. Handling, testing, and conditioning occurred during the dark phase of the light/dark cycle under dim lights (3 lux). During the pre- and post-tests, mice were allowed to explore both chambers freely for 15 min and the time spent in each side was recorded using the ANY-maze behavioral tracking software. We utilized a biased approach in which the unconditioned stimulus was paired with the initially unpreferred side (i.e. the compartment in which the mouse spent less than 50% of the time during pre-test). Mice that displayed a strong initial side preference (i.e. spent ≥65% time in one side) were retired from the experiment prior to conditioning. After the pre-test, mice were randomly assigned to receive i.p. injections of either saline or 20 mg/kg LY466195, followed by an i.p. injection, 15 min later, of either saline or 1.5 g/kg (20% v/v) ethanol during the CS+ conditioning sessions. Conditioning sessions lasted 5 min, with mice receiving the assigned CS+ treatment (CS+ days) or saline-only injections (CS− days) before being placed in the corresponding compartment. Conditioning sessions occurred once daily for eight consecutive days with CS+ sessions every other day. To ensure that mice were actively undergoing withdrawal during testing, subjects were conditioned 20 h after the last ethanol exposure and were given access to 20% ethanol for 4 h after each conditioning session (1:00 PM–5:00 PM). All values are reported as time difference in CS+ chamber (CPP Score).

### 2.8. In vivo Microdialysis

*In vivo* microdialysis was used to measure ethanol-related changes in accumbal DA levels in ethanol-naïve mice and mice with stable consumption in the I2BC drinking paradigm. Mice were stereotactically implanted with a guide cannula (Synaptech Inc, Marquette, MI, USA) within the NAc under 2% isoflurane anesthesia. Surgical asepsis was ensured by cleaning the epilated head with iodine and ethanol prior to making the incision. The following coordinates from Bregma were utilized: A/P +1.5, M/L ±0.9, D/V −4.7. M. The cannula and the anchoring screw were secured with dental cement. Animals were allowed a minimum of 4 days to recover, during which they received a daily administration of 1 mg/kg meloxicam (i.p.) and continued to receive ethanol in the I2BC. Microdialysis probes with a 1-mm membrane (20,000 Da Cutoff, 0.51 mm OD, Synaptech Inc) were inserted into the cannula guide on the afternoon before the microdialysis procedure, and artificial cerebrospinal fluid (aCSF, 149 mM NaCl, 2.8 mM KCl, 1.2 mM CaCl_2_, 1.2 mM MgCl_2_, 0.25 mM ascorbic acid, and 5.4 mM D-Glucose) was infused at a flow rate of 0.2 μL/min overnight. The following day, the aCSF infusion rate was adjusted to 1.1 μL/min for at least 1 h prior to sampling. Samples were collected every 20 min, immediately placed in dry ice and stored at −80° C until analyzed. After collecting four baseline samples, mice were injected i.p. with either saline or 20 mg/kg LY466195. Fifteen min later, mice were injected i.p. with either saline or 2 g/kg ethanol, and six additional samples were collected. At the end of the experiment, mice were perfused, and brains were collected for cannula placement verification. Depending on the experimental group, mice were either maintained on ethanol throughout the testing procedure (ethanol satiety) or had the ethanol removed prior to probe placement (15-20 h ethanol withdrawal).

Dialysis samples were analyzed for DA content by high-performance liquid chromatography using reversed-phase chromatography and electrochemical detection (Dong et al., 2013; Zhang et al., 2012). The system consists of a Thermo Scientific (Waltham, MA) pump (Model 582), an autosampler (Model 542), an HR-80 × 3.2 mm column (3-μm particle size) and a coulometric cell (Model 5014B) connected to an ESA Coulochem II detector. The mobile phase consisted of 4 mM citric acid, 3.3 mM sodium dodecyl sulfate, 100 mM sodium phosphate monobasic monohydrate, 0.25 mM ethylenediaminetetraacetic acid (EDTA), 15% acetonitrile, and 5% methanol. The autosampler mixed 20 μl of the dialysate with ascorbate oxidase (EC 1.10.3.3; 162 units/mg) prior to injection. Quantification of the DA concentration ([DA]) was carried by comparing the peak area relative to external DA standards (0-5 nM). [DA] for each mouse was normalized to the baseline (average of samples 1–4) and represented as % of baseline [DA].

### 2.9. Data and Statistical Analyses

The data are reported as the mean ± standard error of the mean (SEM). SPSS v25 was used to perform linear mixed models analyses with Satterthwaite degrees of freedom. All models included sex as a covariate, and a random intercept for mouse. Statistically significant main effects for drug dose level were followed by Bonferroni adjusted post-hoc tests where each dose level was compared to saline. Statistically significant interaction effects involving drug were followed by linear contrast tests to determine the reason for the interaction.

## 3. RESULTS

### 3.1. LY466195 Modulates Ethanol Consumption and Preference without Altering Sucrose Intake

Ethanol exposure in the I2BC paradigm led to stable, voluntary ethanol intake and preference (Figure 1A & D). The average baseline consumption was calculated using data from weeks 5–6. As previously reported (Hwa et al., 2011), female mice displayed higher baseline intake and preference than males at both time points (Consumption: F_2h-sex_[1,44]=8.66, p<0.01; F_24h-sex_[1,51]=42.57, p<0.001; Preference: F_2h-sex_[1,50]=6.29, p<0.05; F_24h-sex_[1,50]=6.67, p<0.05; data not shown). LY466195 modulated ethanol drinking behavior in a dose-dependent manner, influencing ethanol consumption and preference after 2 h and 24 h of ethanol access (Figure 1B, C, E & F). Administration of 10 mg/kg LY466195 reduced ethanol intake only at the 2-h time point (Figure 1B), whereas 4 mg/kg LY466195 had no effect on drinking phenotype (Figure 1B, C, E & F). Overall, LY466195 did not alter total fluid intake (F[4,144]=1.46, p>0.05), although, there was a difference at 24 h (p<0.01) due to a fluid intake reduction at the 40 mg/kg dose compared to saline (Fig. S1). We also tested whether the effect of LY466195 was moderated by sex but failed to detect a significant interaction at both 2 h and 24 h.

**Figure 1.**
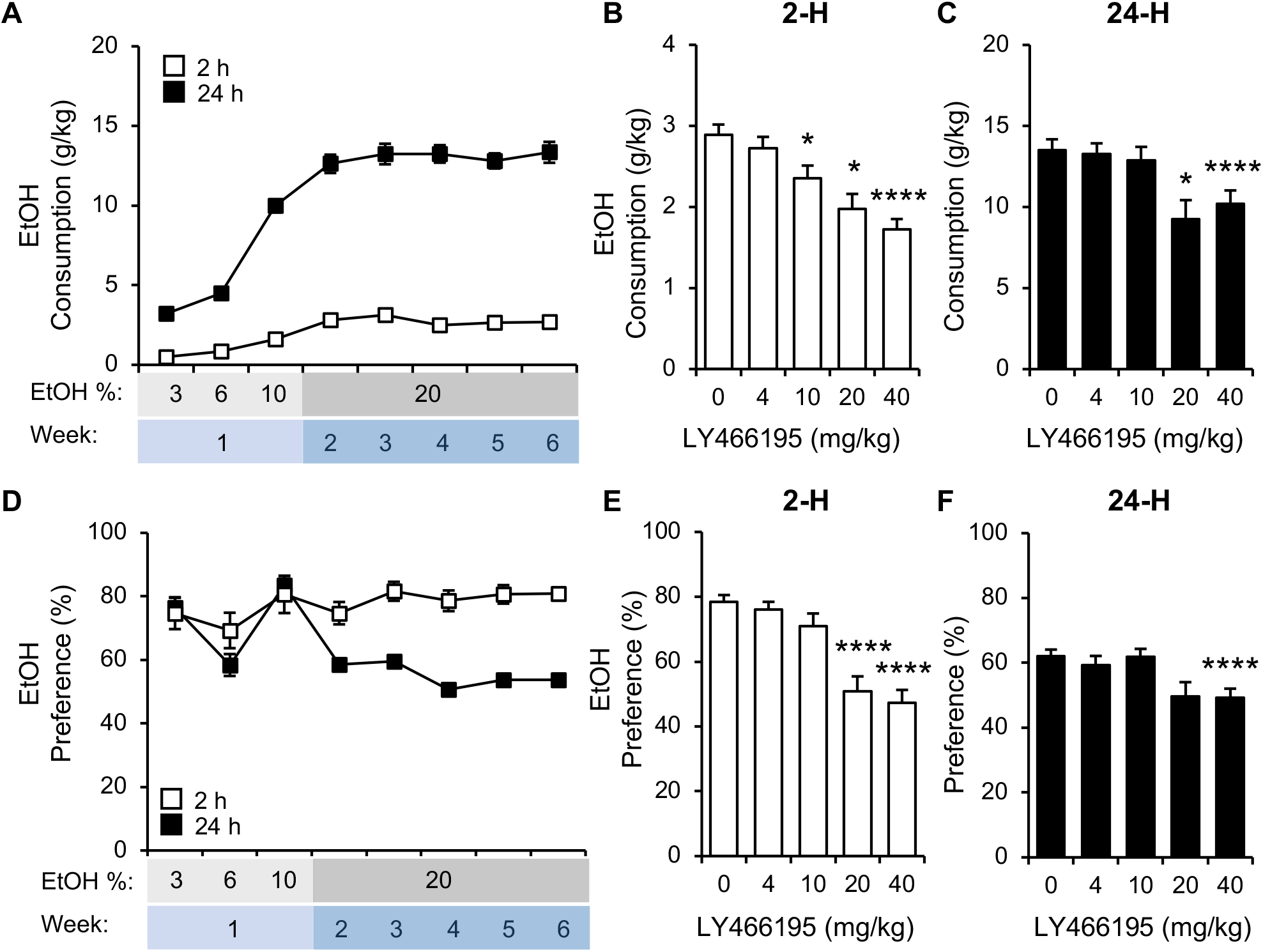
Selective GluK1*KAR blockade modulates ethanol drinking behavior in the I2BC. Mice exposed to the I2BC paradigm for a minimum of 6 weeks display stable ethanol consumption (A) and preference (D). LY466195 was administered via i.p. injections immediately before an ethanol-drinking session. Ethanol dose (B, C) and preference (E, F) at 2 h (B, E) and 24 h (C, F) were modulated by selective GluK1*KAR inhibition in a dose-dependent manner. None of the LY466195 doses examined in this study affected total fluid intake in this drinking paradigm (Supplemental Figure S1). Animal numbers are 56 for A and D, and 56, 47, 39, 20, 48 for B, C, E, and F, respectively. *p<0.05, ****p<0.0001 compared to 0 mg/kg LY466195.

Because LY466195 reduced ethanol consumption after the short, intermittent withdrawal periods of the I2BC paradigm, we proceeded to evaluate the effect of 20 mg/kg LY466195 on ethanol drinking after protracted (6 day-long) withdrawal. We chose this dose because it was the lowest dose to reduce ethanol intake and preference after the short-term withdrawal in the I2BC paradigm (Figure 1), without affecting total fluid intake. We found a significant interaction between LY466195 treatment and repeated withdrawal cycles for both ethanol consumption and preference (Figure 2), with no effect on total fluid consumption. At week 3, LY466195 significantly decreased 2-h ethanol consumption by ~36% when compared to saline-treated animals (p<0.01) and ~40% compared to baseline consumption (p<0.05) (Figure 2A). Administration of LY466195 also attenuated 24-h ethanol consumption but the results did not reach statistical significance during any of the weeks examined (Figure 2B). A significant trend of reduced ethanol preference at both 2-h and 24-h ethanol was observed throughout the testing period in mice treated with LY466195 (Figure 2C, D).

**Figure 2.**
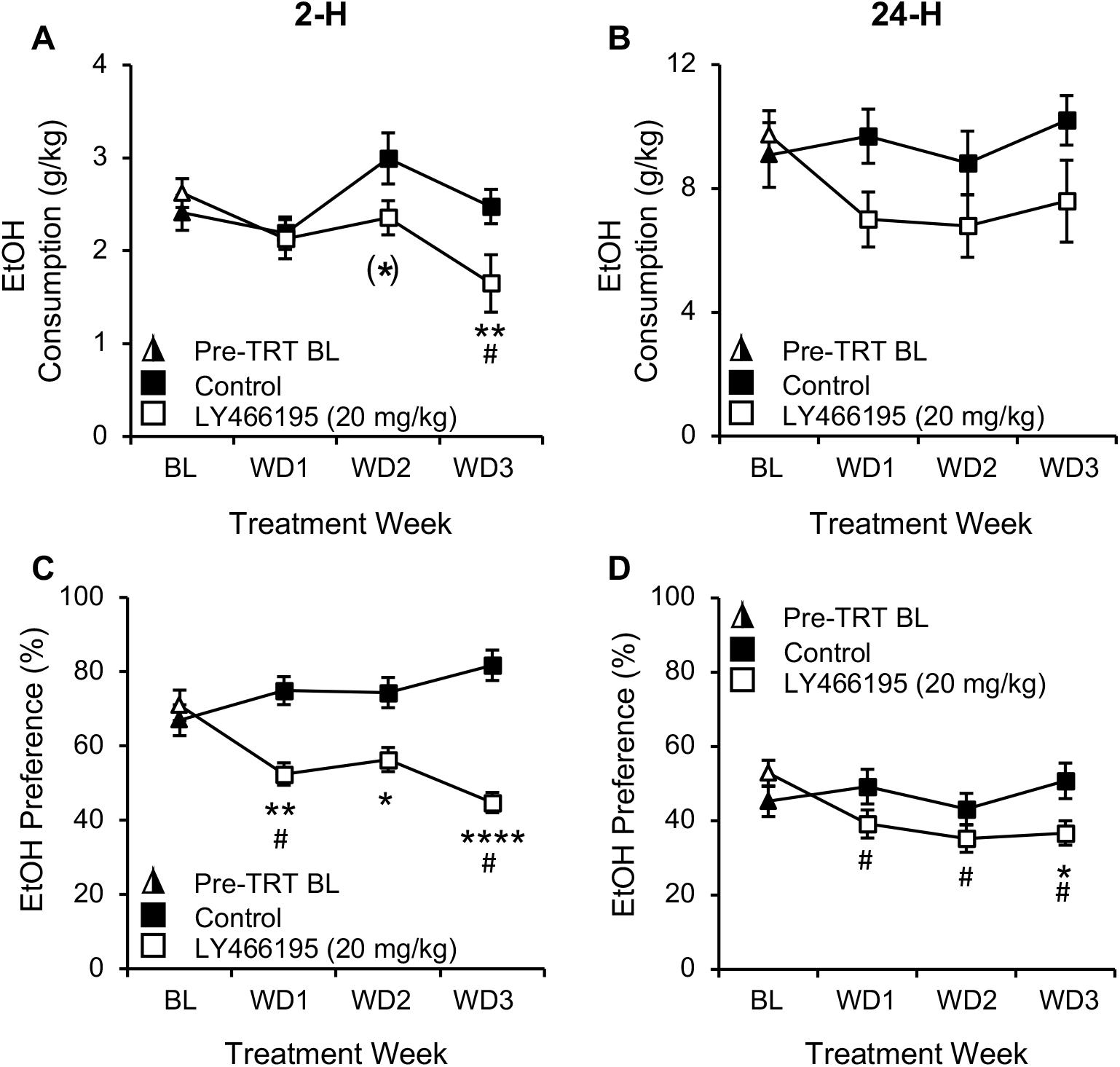
Selective GluK1*KAR inhibition affects ethanol drinking behavior during protracted withdrawal. Mice with stable drinking in the I2BC were exposed to three cycles of protracted withdrawal. For each cycle, the animals were ethanol-deprived for 6 days before a 24-h ethanol re-exposure. An i.p. injection of 20 mg/kg LY466195 was administered immediately before ethanol exposure. Ethanol intake (A,B) and preference (C, D) were determined after 2 h (A, C) and 24 h (B, D) of exposure. Overall, LY466195-treated mice displayed improved drinking behavior. A dose of 20 mg/kg LY466195 significantly reduced 2-h ethanol intake (Week 3) and preference (Weeks 1, 2 and 3) in a cycle-dependent manner. The ethanol preference at the 24-h time point was also significantly decreased in the LY466195-treated group across all weeks when compared to BL. Total fluid consumption was not affected (F_2h_[3,90]=1.64, p>0.05; F_24h_[3,90]=0.62, p>0.05; data not shown). Animal numbers are 17, 15; (*)p=0.05, *p<0.05, **p<0.01, ****p<0.0001 compared to control-treated mice, #p<0.05 compared to BL. TRT = treatment, BL=baseline, WD=ethanol withdrawal. Triangles (▲) represent average BL behaviors in the absence of LY466195.

To determine whether GluK1*KAR inhibition affects natural reward in addition to ethanol drinking, we examined the effect of LY466195 on sucrose intake using a similar in-house drinking protocol (see Supplementary Materials and Methods, Figure S2). Sucrose consumption and preference were unaffected by the acute administration of a 20 mg/kg dose of LY466195 (Supplemental Figures S2A & S2C) while 40 mg/kg LY466195 had an effect on 24-h sucrose consumption (Supplemental Figure S2B, p<0.01). LY466195 at 40 mg/kg had no effect on 24-h preference (Supplemental Figure S2D), but it did affect total fluid intake (F_Total Fluid_[4,82]=5.46, p<0.01). None of the other LY466195 doses we tested affected total fluid intake in this paradigm. Finally, we examined changes in taste preference using a sweet, non-caloric saccharin solution and bitter quinine solutions during treatment with LY466195, as shown in Supplemental Figures S3 and S4, respectively. Overall, we found no significant effects of 20 mg/kg LY466195 on either tastant consumption or preference.

### 3.2. Acute LY466195 Affects Physical Signs of Withdrawal in an Ethanol State-dependent Manner

To test whether GluK1*KAR inhibition modulates the negative reinforcing properties of ethanol, we examined the somatic behavior of ethanol–naïve, –satiated and –withdrawn mice before and after receiving either saline or 20 mg/kg LY466195 (Figure 3). At baseline, in the absence of LY466195 treatment (Figure 3A), withdrawal was characterized by an increase in physical signs. A two-way ANOVA revealed significant main effect of ethanol state and ethanol state by withdrawal sign interaction (F[2,175)=47.40, p<0.0001 and F[8,175)=10.49, p<0.0001, respectively). Post-hoc analysis revealed that mice undergoing withdrawal displayed significantly more shakes, scratches, grooming and “other signs” compared to alcohol-sated mice. The same result was obtained when ethanol-treated mice were compared to ethanol-naïve mice, where withdrawal increased significantly shakes, grooming and “other” signs. Interestingly, we did not detect differences in chewing behavior between the groups. LY466195 treatment affected physical signs with the direction of the effect depending on the alcohol state. A two-way test of the physical behaviors post-LY466195 administration revealed a significant ethanol-state by withdrawal sign interaction (Figure 3B; F[8,90]=3.731,p<0.001). Following LY466195 treatment, a significant difference between naïve and withdrawal group was detected for grooming (p<0.05). Interestingly, we observed significantly higher levels of chewing in the satiated group compared to both naïve (p<0.01) and withdrawal (p<0.001) groups. The effect of LY466195 treatment was also examined in each group (i.e. naïve, satiety and withdrawal) compared to a test following a saline i.p. injection. LY466195 had no effect in ethanol naïve mice (Figure 3C; acute treatment: F[1,40]=0.17, p=0.6863); acute treatment by withdrawal sign: F[4,40]=1.325, p=0.2775). In alcohol satiated mice (Figure 3D), we detected a significant main effect of acute treatment (F[1,65]=5.483, p<0.05, and acute treatment by withdrawal sign interaction (F[4,65]=5.819, p<0.001). As already detected, the GluK1R antagonist significantly increased chewing behavior (saline vs. LY466195 alcohol-satiated mice: p<0.0001) but had no significant effect on any of the other categories of physical signs. Lastly, for mice undergoing withdrawal (Figure 3E), a two-way ANOVA also detected a significant main effect of acute treatment (F[1,55]=11.51, p<0.005) and acute treatment by sign interaction (F[4,55)=5.908, p<0.001). Post-hoc analysis revealed that LY466195 significantly reduced withdrawal-induced shakes (p=0.0010) while other signs of withdrawal were not altered.

**Figure 3.**
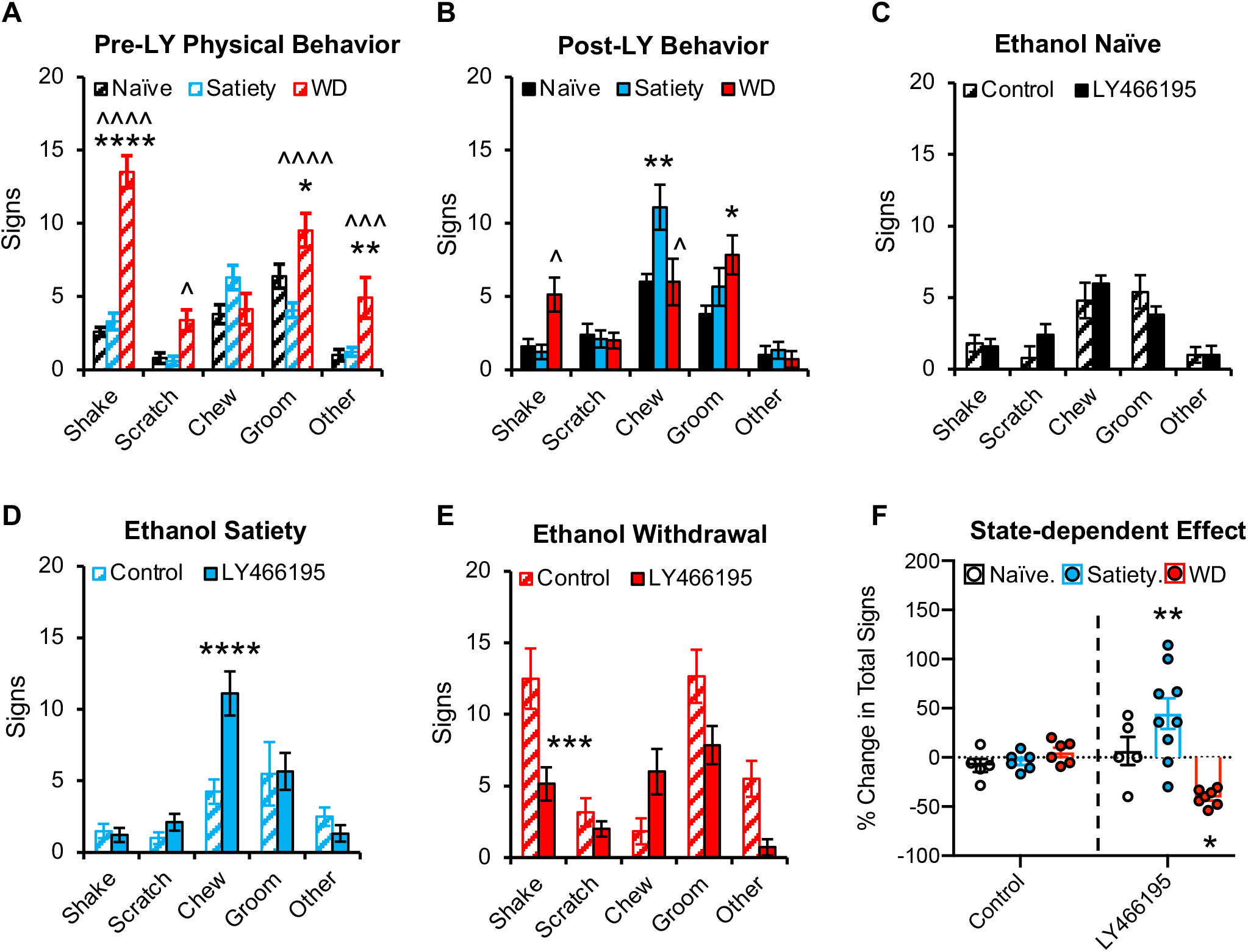
GluK1*KAR blockade has opposite effects on physical signs, depending on the ethanol state. Various physical behaviors were evaluated in ethanol-naïve, -satiated, and -withdrawn mice before (A) and following an i.p. injection of 20 mg/kg LY466195 (B). Mice undergoing spontaneous 24-h withdrawal displayed significantly more physical signs of withdrawal during the pre-test, compared to ethanol -naïve and -satiated mice (A). An acute administration of LY466195 decreased the occurrence of various signs of withdrawal in ethanol-withdrawn mice but increased the occurrence of chewing in the satiated group (B). An injection of 0.01 mL/g saline was used as a control for each ethanol state group (C, D, E), which did not significantly affect the manifestation of physical behaviors as evidenced by a negligible percent change in total number of signs (F). A comparison of the percent change observed in each group (F) revealed that the effect of LY466195 on somatic signs was dependent on the ethanol state. The GluK1*KAR inhibitor had no effect on ethanol-naïve mice (C), whereas the total number of signs was significantly changed in ethanol-treated mice (D, E). Ethanol-withdrawn mice displayed a significant decrease, whereas ethanol-satiated mice showed an increase in total score. For A, n’s are 10, 14, 13. For B, n’s are 5, 9, 7. For C, n’s are 5, 5. For D, n’s are 6 and 9. For E, n’s are 6 and 7. For F, n’s are 5, 5, 6, 9, 6, 7. For A-B, *p<0.05, **p<0.01, ****p<0.0001 compared to Naïve, and ^p<0.05, ^^p<0.01, ^^^^p<0.0001 compared to Satiety. For C-F, *p<0.05, **p<0.01, ***p<0.001 compared to correspondent saline-treated group.

To further illustrate the interaction between ethanol state and LY466195 treatment, Figure 3F shows the percentage change from baseline in total number of physical signs. A two-way ANOVA of these data detected a significant main effect of ethanol state (F[2,32]=6.554, p<0.01) and ethanol state by acute treatment interaction (F[2,32)=10.21, p<0.001). Post-hoc analysis of the percent changes revealed a significant increase in satiety (p<0.01), but a significant decrease in withdrawal (p<0.05). In summary, these results highlight the opposite effect exerted by LY466195 in satiated vs. withdrawn mice, suggesting that GluK1*KAR function differs depending on the ethanol state.

### 3.3. GluK1*KARs Modulate Ethanol CPP during Periods of Ethanol Cessation

To investigate whether LY466195 reduces ethanol consumption by altering ethanol’s rewarding properties, we assessed the drug’s effect on the acquisition of ethanol CPP. A total of 8 mice (4 each from the ethanol naïve and ethanol-treated groups) were removed from the experiment due to initial side preference (≥65%) in the pre-test. LY466195 had no effect on the development of ethanol CPP in ethanol-naïve animals nor did it have any effect in control, saline-treated mice (Figure 4A). In ethanol-naïve mice, a two-way test revealed a main effect of ethanol (F_EtOH_[1,60]=6.41, p<0.05), but not LY466195 (F_LY_[1,60]=0.19, p>0.05) or a drug x ethanol state interaction (F[1,60]=0.07, p>0.05) (Figure 4A). Given that we observed a state-dependent effect on the physical manifestations of withdrawal, we repeated the experiment in mice undergoing ethanol withdrawal (Figure 4B). In those animals, the two-way test revealed a drug x ethanol state interaction (F[1,30]=5.38, p<0.05), such that mice undergoing spontaneous withdrawal did not display ethanol CPP when conditioned using a dose of 1.5 g/kg EtOH (Figure 4B). Conversely, LY466195 treatment prior to ethanol administration led to greater ethanol CPP in ethanol–withdrawn mice (p<0.05). Importantly, mice treated with LY466195 in the absence of ethanol did not display CPP, suggesting that the drug, even in a negative state, is not rewarding on its own.

**Figure 4.**
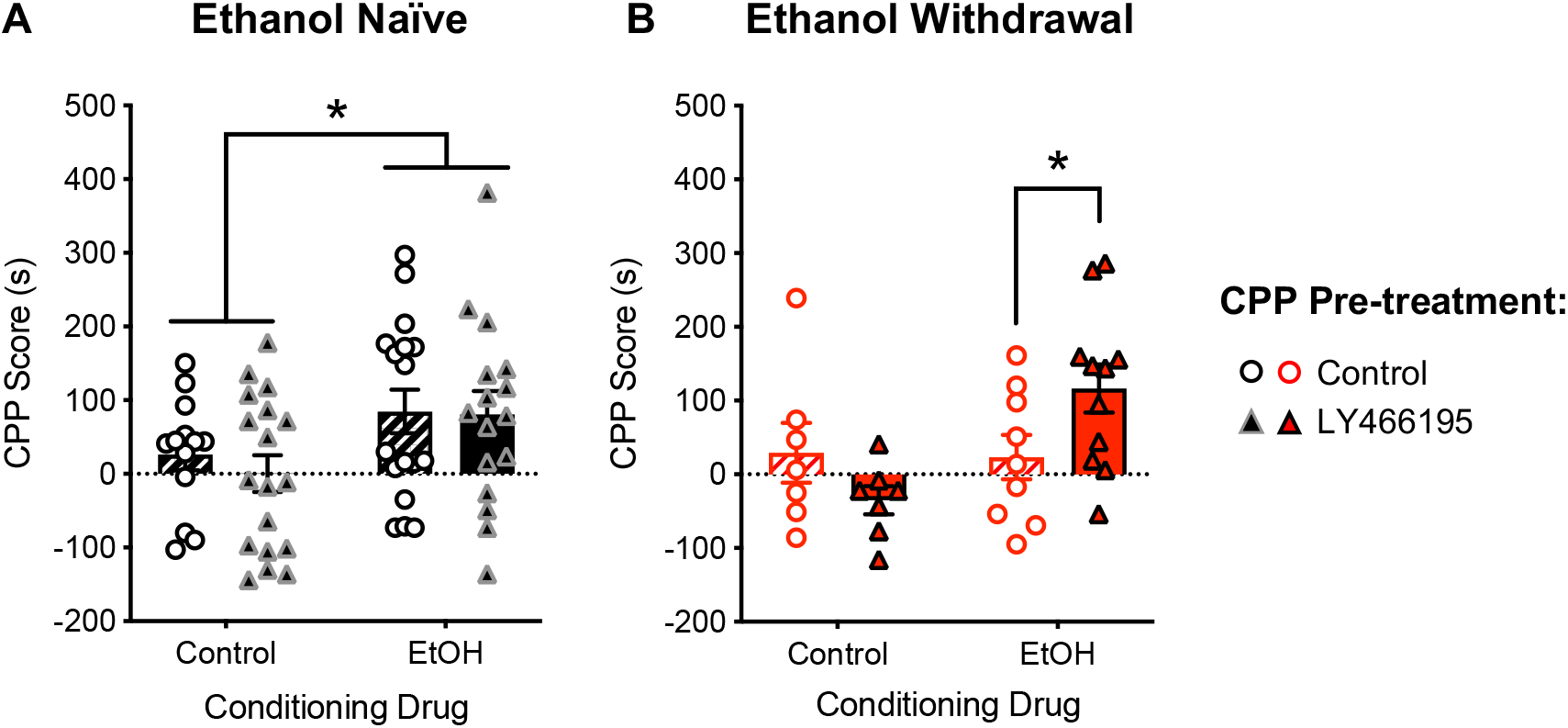
Ethanol withdrawal prevents the acquisition of ethanol CPP, a behavior that is rescued by selective GluK1*KAR blockade. Ethanol-induced CPP was evaluated in ethanol –naïve (A) and ethanol – withdrawn (B) mice following an injection of either saline or 20 mg/kg LY466195. Ethanol-naïve mice displayed strong ethanol CPP, whereas this behavior was not observed in mice undergoing spontaneous ethanol withdrawal. An acute administration of LY466195 did not affect the development of ethanol CPP in ethanol–naïve mice but rescued the behavior in withdrawn mice. CPP was not observed in mice treated with LY466195 but not ethanol, suggesting that this drug is not rewarding on its own. Animal numbers are 13, 18, 17, 16 for ethanol-naïve, and 7, 7, 9, 11 for ethanol withdrawal; *p<0.05 compared to indicated group/s.

### 3.4. GluK1*KARs Modulate Ethanol-related Changes in DA Levels in the NAc during Ethanol Withdrawal

To investigate whether treatment with 20 mg/kg LY466195 alters ethanol-induced changes in accumbal dopamine concentrations ([DA]) in an ethanol state-dependent manner, ethanol-naïve, -satiated, and -withdrawn mice received one of two treatments: saline-ethanol or LY466195-ethanol. A third treatment paradigm, LY466195-saline, was used to examine the intrinsic ability of LY466195 to modify dopaminergic activity. Baseline DA levels in the three ethanol states did not differ significantly (Figure 5B; F[2,43]=2.132, p=0.1310). To investigate the effect of acute co-treatment on levels of DA in the NAc, we examined the effect of injections on percent basal [DA] using a three-way ANOVA with repeated measures (RM), which detected a significant co-treatment-by-state-by-time interaction (F[19,177]=4.148, p<10^−6^). To further interpret the three-way interaction, we conducted a series of RM two-way ANOVAs to examine the effect of ethanol state at different levels of acute co-treatment. We also computed simple main effects using RM two-way ANOVAs for the effect of acute co-treatment at different levels of ethanol state. RM two-way ANOVA for the saline-ethanol treatment revealed significant main effects of ethanol state (F[2,14]=4.405, p<0.05) and time (F[4.8,66.9]=13.68, p<0.0001), and a significant state-by-time interaction (F[18,126]=2.129, p=0.01). Upon closer examination of the temporal effects of acute treatment and ethanol state on accumbal DA levels, we first observed that in mice receiving the saline-ethanol treatment, the ethanol state influenced the time course to return to basal DA levels without affecting peak of percent basal [DA]. Specifically, while there was no significant difference between dopaminergic responses in naïve and satiated mice (Supplemental Figure S5A), there was a significant difference between ethanol-naïve and ethanol-withdrawn mice (Figure 5C), with ethanol injection leading to significantly higher DA levels for ~ 60 minutes (minutes 155 to 215) post-ethanol injection in mice undergoing withdrawal (155min: p=0.010; 175min: p<0.01; 195min: p<0.05). We then examined data for LY-SAL using a RW two-way ANOVA, which also revealed significant main effects of state (F[2,11]=4.954; p<0.05) and time (F[2.8,30.7]=7.122, p<0.01), and a significant state-by-time interaction (F[18,99]=2.525, p=0.01). We did not detect significant differences between naïve and satiety following LY-SAL treatment (Supplemental figure S5C). However, LY466195 injection in mice undergoing withdrawal produced significantly higher DA levels compared to naïve mice (Figure 5D; 135min: p<0.05; 155min: p<0.05), suggesting that the GluK1 antagonist can modify dopaminergic activity depending on the ethanol-state. We finally wanted to examine whether treating the mice with LY466195 before acute ethanol injection (i.e. LY-ETOH) had an effect on accumbal DA levels for each level of the state variable. For ethanol naïve mice (Figure 5E), a RM two-way ANOVA revealed significant main effects of time (F[4, 40]=10.94, p<0.0001) and pre-treatment (F[1,10]=10.02, p<0.05), and a significant time-by-pre-treatment interaction (F[9,90]=9.428, p<0.0001). Post-hoc analysis revealed that LY-ETOH treated, naïve mice had higher levels of percent basal DA than those treated with SAL-ETOH. Interestingly, although we detected significant time-by-pre-treatment interaction (F[9,81]=3.240, p<0.01) in mice undergoing withdrawal (Figure 5F), we failed to detect significant differences using the Bonferroni’s multiple comparison’s test. Importantly, the responses between LY466195-ethanol-treated mice undergoing withdrawal (Figure 5F, triangles) and naïve mice treated with saline-ethanol were comparable (Figure 5E, triangles). Finally, LY466195 injection prior to ethanol did not result in overall significant differences between naïve and alcohol satiated mice (F_state_[1,7]=1.761, p=0.2262; F_state-by-time_[9,63]=1.587, p=0.1385; Supplemental Figure S5B).

**Figure 5.**
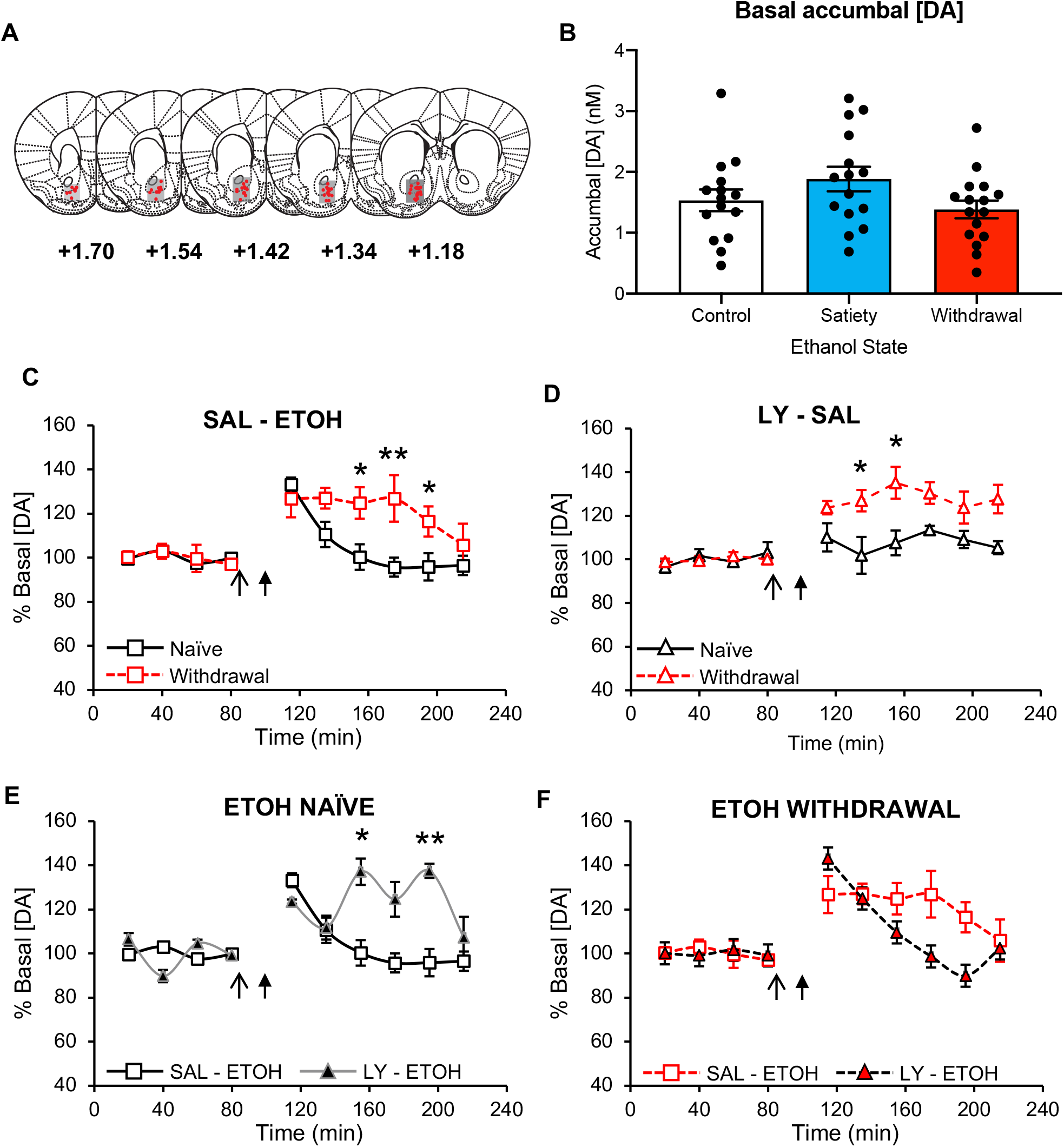
Acute ethanol-related accumbal DA overflow is modulated by selective GluK1*KAR blockade. DA levels in the NAc were measured in ethanol –naïve and –withdrawn mice following an injection of either saline or 20 mg/kg LY466195 (open arrow) that was followed by either an ethanol or saline injection (closed arrow). (A) Anatomical distribution of dialysate sampling areas in the NAc. The opacity of the gray squares is indicative of the percentage of mice with probes located on that anterior-posterior (AP) plane, and the red dots represent the tip of the dialysate probes (1-mm membrane). (B) Baseline DA levels were recorded for 80 min prior to acute treatment in ethanol naïve, -satiated, and withdrawn mice. (C-F) Ethanol-evoked increase in [DA] was influenced by ethanol state and acute treatment. The response to ethanol was significantly prolonged in ethanol-withdrawn mice compared to ethanol-naïve mice following an acute ethanol administration (C). In ethanol-withdrawn mice, the co-administration of LY466195 and ethanol produced similar [DA] as those observed in ethanol-naïve animals that received saline + ethanol (E and F). Testing the effect of LY466195 in the absence of ethanol suggests that GluK1*KAR antagonism promotes an increase in accumbal [DA] in ethanol-withdrawn, but not in ethanol-naïve mice (D). For (B), animal numbers are 15, 15 and 16 for naïve, satiety and withdrawal, respectively. For (C-F), animal numbers are 8 (naïve, SAL-ETOH), 5 (withdrawal, SAL-ETOH), 4 (naïve, LY – ETOH), and 5 (withdrawal, LY – ETOH), 3 (naïve, LY-SAL), and 5 (withdrawal, LY-SAL). *p<0.05, **p<0.01, ***p<0.001 compared to control at the specified time-point. ETOH = ethanol, LY = LY466195 and SAL = Saline.

## 4. DISCUSSION

We examined the role of a GluK1*KAR inhibitor, LY466195, in reducing ethanol consumption in a mouse model of chronic, voluntary intake. The work was based on ethanol’s effects on excitatory signaling at ionotropic glutamate receptors (Smart and Paoletti, 2012), including the kainate subtype (Rao et al., 2015), and evidence that TPM’s effects on drinking are moderated by variation in *GRIK1* (Kranzler et al., 2014a), the gene that encodes the GluK1 subunit of the KAR. We found that GluK1*KAR blockade can influence several drinking-related behaviors. First, LY466195 reduced ethanol consumption and preference in the I2BC paradigm. These results are in accordance with a recent study showing decreased ethanol preference in the I2BC paradigm in Long Evans rats treated with LY466195 (Van Nest et al., 2017). In that study, the decrease in ethanol preference was accompanied by an increase in total fluid intake, which we did not observe in mice. Although in rats there was no effect of LY466195 on ethanol intake, the highest dose of LY466195 examined was 10 mg/kg, which in mice reduced ethanol intake only during the first 2 hours of ethanol exposure (Figure 1B). Thus, higher doses of the drug (i.e., 20 or 40 mg/kg LY466195) might have also reduced ethanol intake and preference in rats. A potential explanation for the reduced ethanol consumption would be a change in ethanol metabolism. However, in mice, *Grik1*, as well as *Grik3* and *Grik5* (the genes that encode the other KAR subunits potentially targeted by LY466195) have very low to no expression in the liver (GenBank accession number: PRJNA66167), where ethanol is primarily metabolized. Thus, we do not predict LY466195 to have a significant effect on ethanol metabolism.

LY466195 reduced ethanol intake and preference after prolonged periods of withdrawal (Figure 2), suggesting that GluK1*KAR antagonists could help to prevent drinking relapse. TPM is also effective in reducing alcohol intake and preventing relapse in humans (Baltieri et al., 2008; Florez et al., 2008). Therefore, although TPM inhibits both AMPA and kainate receptors (Gryder and Rogawski, 2003; Skradski and White, 2000), our data point to an important role of GluK1*KAR antagonism in the therapeutic effects of TPM. In addition to reducing ethanol intake and preference during withdrawal, LY466195 also reduced the physical signs of withdrawal (Figure 3C and 3F). Following long-term ethanol consumption, the brain adapts to the depressant effects of ethanol by increasing excitatory (e.g., glutamatergic) and reducing inhibitory (e.g., GABAergic) neurotransmission (Bell et al., 2016; Rao et al., 2015). Elevated brain glutamate levels associated with these homeostatic mechanisms could contribute to ethanol-withdrawal-related negative states (Läck et al., 2009, 2008). Thus, excessive stimulation of GluK1*KARs could contribute to craving and severe manifestations of withdrawal, which could be reduced by GluK1*KAR antagonism.

We were unable to examine the effects of LY466195 on affective signs of withdrawal because this drug altered locomotor behavior in ethanol-naïve mice tested in the open field arena and the elevated plus maze (Supplemental Figure S6). Altered locomotion can affect the results of anxiety tests that rely on the animal moving to explore the testing apparatus. Hyperlocomotion following LY466195 treatment in rats was not reported by Van Nest and colleagues (2017), possibly reflecting species-specific differences. It should also be noted that LY466195-treated mice exhibited neither hyperlocomotion nor any other abnormal behavior when observed in the home cage.

Ethanol enhances the activity of the dopaminergic system, resulting in increased DA levels in the NAc (Chiara and Imperato, 1988; Gessa et al., 1985; Imperato and Di Chiara, 1986), a key mediator of ethanol reward. We observed that LY466195 did not affect ethanol CPP in ethanol-naïve mice (Figure 4). We also found that mice chronically treated with ethanol in the I2BC paradigm and undergoing withdrawal during conditioning did not develop CPP to 1.5 g/kg EtOH, which elicits a strong CPP in ethanol-naïve mice (Figure 4). To our knowledge, this is the first study to assess the effect of spontaneous ethanol withdrawal following the I2BC paradigm on ethanol CPP in response to a systemic ethanol dose shown by us and others to produce CPP in non-dependent mice (Groblewski et al., 2008). A previous study examined the effect of chronic ethanol exposure on ethanol CPP after 4 days of ethanol liquid diet using 24 conditioning sessions and various doses of ethanol (Ting-A-Kee et al., 2009), where it was reported that ethanol-treated mice developed CPP at 0.2 g/kg but not at higher doses (such as 0.8 and 2.0 g/kg). Such findings suggest that there might be a leftward shift in the dose-response curve after chronic ethanol treatment in mice. Here we show that 1.5 g/kg EtOH was insufficient to produce CPP during withdrawal following chronic ethanol exposure. Interestingly, mice conditioned with LY466195 and ethanol during withdrawal developed CPP (Figure 4), suggesting that LY466195 treatment can compensate for some of the neuroadaptations associated with chronic ethanol exposure. In the future, examining the effect of LY466195 treatment on CPP to lower ethanol doses during withdrawal would clarify if inhibiting the GluK1*KARs is shifting the dose-response curve back to non-dependent levels and correcting for sensitization.

Overall, our behavioral experiments point to a state-dependent effect of GluK1*KARs in modulating ethanol-associated behaviors. DA changes within the NAc have been documented during different phases of the alcohol addiction process, with acute alcohol increasing and chronic alcohol leading to reduced DA levels (Gilpin and Koob, 2008). Therefore, we investigated whether LY466195 treatment altered the effect of acute ethanol on DA levels in the NAc in an ethanol state-dependent manner. We found the effects of acute ethanol exposure on accumbal DA levels to differ depending on ethanol state (Figures 5C and Supplemental Figure S5A). While acute ethanol produced DA increases that were similar in size and time course between ethanol-naïve and ethanol satiated mice, the same acute ethanol exposure led to a prolonged increase in accumbal DA overflow during withdrawal (Figure 5C). Previous studies have suggested that the biphasic dopaminergic response to ethanol on DA release in the NAc is due to differential involvement of receptor systems and brain regions after an ethanol administration. Particularly, GABA_A_ receptors expressed in the NAc have been shown to play a crucial role in DA decline from peak values after ethanol exposure (Löf et al., 2007; Söderpalm et al., 2009). Therefore, the prolonged response observed in ethanol-withdrawn mice might be at least partially explained by changes in GABA_A_ receptor responses to ethanol in the NAc. However, it was very interesting to observe that when acute ethanol exposure was preceded by an injection of LY466195 in mice undergoing withdrawal, the peak DA level and the time to return to baseline were indistinguishable from those in ethanol-naïve animals treated with saline-ethanol (Figure 5E-F), suggesting that the dynamics of ethanol-induced DA release during withdrawal are influenced by GluK1*KARs antagonism. Specifically, this observation suggests that GluK1*KARs might also contribute to the time course of dopamine release after chronic ethanol exposure, although further studies using genetic approaches are needed to explore this hypothesis and discard the possibility that our observation is the result of off-target effects of LY466195.

We also observed a state-dependent effect of LY466195 on the manifestation of withdrawal, with increased physical signs in ethanol-satiated animals and decreased signs in ethanol-withdrawn mice (Figure 3). Based on this behavioral effect and its state-dependent influence on the dopaminergic system, we hypothesize that LY466195 could be administered to subjects that are current alcohol abusers to produce alcohol aversion, and to subjects that are abstinent to mitigate the behavioral and neurochemical correlates of withdrawal. Overall, our observations suggest that the GluK1*KAR system changes in response to chronic ethanol treatment. Whereas previous studies found no changes *in GRIK1* expression in the brain following chronic ethanol treatment (Jin et al., 2014a, 2014b), the GluK1 subunit could be integrated in KARs of different subunit composition or the GluK1*KAR signaling mode (i.e., ionotropic vs. metabotropic) could change during chronic ethanol exposure and withdrawal (Lerma and Marques, 2013). Our findings also suggest that a delicate balance exists in the regulation of GluK1*KAR-mediated transmission, with deviations from baseline in either direction affecting the system’s response to ethanol. Despite strong evidence that glutamatergic transmission is involved in alcoholism (Gass and Olive, 2008), it remains unclear how KAR-mediated synaptic transmission is altered by chronic ethanol consumption and how the effects may vary among different brain regions.

Compulsive ethanol drinking is associated with aberrant plasticity that occurs in multiple brain regions, including the mesocorticolimbic and extended amygdala reward circuits (Koob, 2013; Nam et al., 2012; Tabakoff and Hoffman, 2013). Because baseline anxiety levels and the anxiety that develops during ethanol withdrawal can contribute to the initiation and maintenance of alcohol abuse (Bibb and Chambless, 1986; Wilson, 1988; Koob, 2003; Cardoso et al., 2006), the specific role of GluK1*KAR in the amygdala also warrants investigation. The amygdala expresses high levels of KARs, including GluK1*KARs, which mediate a form of heterosynaptic plasticity that results in enduring excitatory synaptic responses (Li et al., 2001). This neuroadaptation could also participate in the effects of chronic ethanol exposure on compulsive drinking and the signs of ethanol withdrawal.

## 5. CONCLUSIONS

In summary, we identified an important role of GluK1*KAR-mediated neurotransmission in modulating ethanol consumption and pinpointed the mechanisms that could underlie the efficacy of GluK1*KAR blockade in reducing drinking. GluK1*KARs appear to be key modulators of both the positive and negative reinforcing properties of ethanol, particularly during withdrawal following chronic administration. Future studies should further examine the neuroplasticity that this receptor system undergoes after prolonged ethanol exposure, to possibly design more efficacious and better tolerated medications, such as GluK1*KAR antagonists, that reduce drinking in patients with AUD.

## ABBREVIATIONS

AUD: alcohol use disorder
FDA: US Food and Drug Administration
TPM: topiramate
SNP: single nucleotide polymorphism
KAR: kainate receptor
GluK1*KAR: GluK1-containing kainate receptor
EtOH: ethanol
BLA: basolateral amygdala
NAc: nucleus accumbens
B6: C57Bl/6J
I2BC: intermittent two-bottle choice
i.p.: intraperitoneal
CPP: conditioned place preference
aCSF: artificial cerebrospinal fluid
HPLC: high performance liquid chromatography
DA: dopamine
[DA]: DA concentration

## ACKNOWLEDGMENTS

The authors would like to acknowledge Luke Egan and Shreya Ganguly for their assistance with the tastant consumption and CPP experiments, respectively. The authors would also like to thank Lilly Pharmaceuticals for generously donating LY466195 for use in this study.

