## Supplemental figures for "Antagonism of GluK1-containing Kainate Receptors Reduces Ethanol Consumption by Modulating Ethanol Reward and Withdrawal"

### SUPPLEMENTAL MATERIAL and METHODS

#### Sucrose, Saccharin and Quinine Consumption

To assess the specificity of LY466195, we tested the effect of acute drug treatment on sucrose (caloric reinforcer), saccharin (non-caloric reinforcer), and quinine (bitter, aversive) drinking on single-housed, ethanol-naïve mice using a continuous two-bottle choice protocol, in which one bottle contained filtered water and the other contained the tastant solution. The tastants were presented in the following order: 2% sucrose (w/v), 0.02% saccharin (w/v), and 0.03 mM quinine. The mice had access to each tastant for at least 3 weeks, and the positions of the bottles were changed daily to avoid the acquisition of side preference. During the first week of tastant exposure, the mice were acclimated to the taste, and baseline consumption and preference after 2 h and 24 h of consumption was determined. During the second and third weeks, the mice were acutely treated on Monday, Wednesday and Friday with i.p. injections of either saline, 4, 10, 20, or 40 mg LY466195/kg. Every mouse received each LY466195 treatment, and the order of the doses was randomized. The mice were weighed at least once a week to determine tastant consumption and preference, as described for the I2BC protocol.

#### Acute Effect of LY466195 on Locomotive Activity

The effect of acute, systemic administration of LY466195 was evaluated in adult, ethanol-naïve mice in the open field arena (OFA). Briefly, the OFA consisted of a matte white Plexiglass® acrylic box measuring 40 cm x 40 cm x 20 cm (D x L x H). After a one-week habituation period to daily 0.1 mL/g saline i.p. injections, mice were tested in the OFA for 30 min under dim-light conditions (~3 lux) following an administration of either saline, 4 mg/kg, or 20 mg/kg LY466195. The locomotive behavior of the mice was examined by measures of total distanced traveled in the OFA obtained using the ANY-maze ™ video tracking system. Additionally, to test whether the effect of LY466195 on locomotion was transient, a subset of mice was repeatedly tested in the OFA. For these experiments, mice were maintained in the same treatment group and testing occurred every other day.

**SUPPLEMENTAL FIGURE LEGENDS**

***Figure S1*. Dose-response curve of the effect of acute LY466195 administration on total fluid intake in the I2BC**. Mice with stable drinking in the I2BC were treated with acute doses of LY466195 (i.p.). Total fluid consumption was measured 2-h (white) and 24-h (gray) following LY466195 administration. One-way ANOVA did not detect an effect of acute treatment on total fluid intake ($F[4, 144]=1.458$, $p=0.218$). Nonetheless, a significant effect of acute treatment was found at 24-h ($F[4, 151]=4.065$, $p=0.004$). Bonferroni's multiple comparisons test revealed a significant difference between 0 and 40 mg/kg ($p=0.003$). Importantly, 20 mg/kg LY466195, which is the dose we used throughout the study, did not lead to significant changes in total fluid intake ($p_{2\text{-}h}=0.357$, $p_{24\text{-}h}=0.601$). Animal numbers are 56, 47, 40, 20, 48. *$p<0.05$ compared to 0.

***Figure S2.* Moderate acute doses of LY466195 do not affect voluntary sucrose consumption.** Ethanol-naïve mice with continuous access to 2% sucrose in a two-bottle choice were injected i.p. with either saline or LY466195 at 1:00 PM (3 h into the dark portion of their light cycle). Two-hour (A, C) and twenty-four-hour (B, D) sucrose consumption (A, B) and preference (C, D) were examined in response to selective GluK1*KAR inhibition. A main effect of LY466195 treatment on 24-h sucrose consumption was detected ($F[4,82]=5.48$, $p=0.001$), where 40 mg/kg LY466195 significantly decreased tastant intake compared to control treatment ($p=0.002$). A parallel effect was observed for 24-h total fluid intake ($F[4,82]=5.46$, $p=0.001$), where mice that received 40 mg/kg LY466195 displayed decreased fluid consumption compared to control-treated mice ($p=0.002$). However, 20 mg/kg LY466195 did not modulate sucrose intake or preference, and it did not alter total fluid intake. Animal numbers are 30, 30, 13, 12, 30.

***Figure S3.*** **Acute LY466195 promotes a transient, dose-dependent effect on voluntary saccharin consumption.** Ethanol-naïve mice with continuous access to 0.2% saccharin in a two-bottle choice were injected i.p. with either saline or LY466195 at 1:00 PM (3 h into the dark portion of their light cycle). Two-hour (A, C) and twenty-four-hour (B, D) saccharin consumption (A, B) and preference (C, D) were examined in response to selective GluK1*KAR inhibition. A main effect of LY466195 dose on 2-h saccharin consumption was found (F[4,108]=6.32, p<0.001), where 10 mg/kg increased sucrose intake at this time point (p=0.003). We also observed an main effect of LY4666195 treatment on 2-h total fluid intake (F[4,108]=9.52, p<0.001), where mice that were treated with 10 mg/kg LY466195 had significantly higher 2-h total fluid intake compared to saline treated mice (p=0.008). However, the effect of this dose was transient, with no modulation at the 24-h time point. Importantly, 20 mg/kg LY466195 (the dose that was utilized to assess ethanol-related behaviors) did not alter 2-h and 24-h sucrose intake nor preference. Animal numbers are 28 for each dose.

***Figure S4.*** **Acute LY466195 does not affect voluntary quinine consumption.** Ethanol-naïve C57Bl/6J mice with continuous access to 0.03 mM quinine in a two-bottle choice were injected i.p. with either saline or LY466195 at 1:00 PM (3 h into the dark portion of their light cycle). Two-hour (A, C) and twenty-four-hour (B, D) quinine consumption (A, B) and preference (C, D) were examined in response to selective GluK1*KAR inhibition. Overall, LY466195 did not modulate quinine intake or preference. However, it is important to mention that we detected a main effect of LY466195 treatment on total fluid intake at 2 h (F[4,114]=9.57, p<0.001) and 24 h (F[4,92]=4.93, p=0.001). Post hoc analysis revealed a reduction in total fluid intake only in mice treated with 40 mg/kg LY466195 ($p_{2h}$=0.007, $p_{24h}$<0.001). No effect was detected for 20 mg/kg LY466195, the dose used in most experiments, on any of the parameters that we examined. Animal numbers are 24 for each dose.

***Figure S5.*** **Ethanol-related accumbal DA overflow is similar in naïve and ethanol-satiated mice after selective GluK1*KAR blockade.** DA concentrations in the NAc were measured in ethanol–naïve and satiated mice following an injection of either saline or 20 mg/kg LY466195 (open arrow), and either an acute ethanol or saline injection (closed arrow) fifteen minutes later. DA concentrations in 20-min intervals were normalized to the average baseline concentration for ethanol –naïve, and –satiated mice. Mice were either treated with saline + ethanol (A), LY466195 + ethanol (B), or LY466195 + saline (C). *p<0.05 compared to ethanol naïve mice at the specified time-point. For ethanol naïve, Animal numbers are 8 for SAL-EtOH, 4 for LY-EtOH, and 3 for LY-SAL. For ethanol satiety, Animal numbers are 4 for SAL-EtOH, 5 for LY-EtOH, and 6 for LY-SAL. LY = LY466195, EtOH = ethanol, SAL = saline.

***Figure S6.*** **Acute LY466195 administration increases locomotor activity in the OFA**. Locomotion was examined in ethanol naïve mice in the OFA following an injection of either saline, 4 mg/kg, or 20 mg/kg LY466195. (A) LY466195 resulted in increased total distance traveled in the OFA (F[2, 39]=10.67, p=0.0002). Bonferroni's multiple comparisons test revealed that both 4 mg/kg and 20 mg/kg LY466194 increased total distance traveled in the OFA compared to saline ($p_{4\text{-mg/kg}}$=0.0057, $p_{20\text{-mg/kg}}$=0.0004). Interestingly, we failed to detect an effect of dose on the effect of LY466195, with no statistical difference between the 4-mg/kg and 20-mg/kg groups (p>0.9999). (B) Repeated exposures to LY466195 did not mitigate the effect of the drug on locomotion. A two-way ANOVA revealed a significant main effect of acute treatment on total distance traveled (F[2, 18]=12.70, p=0.0004), but no interaction between testing day and acute treatment (F[4, 36]=1.027 ,p=0.4064). As expected, we observed a significant main effect of testing day on total distance traveled (F[2, 33]=38.05, p<0.0001), related to familiarization to the testing apparatus. Animal numbers are 18, 10, 14 for (A) and 6, 7, 8 for (B). *p<0.05, **p<0.01, ***p<0.001.

**Figure S1.** Dose-response curve for the effect of acute LY466195 administration on total fluid intake in the I2BC

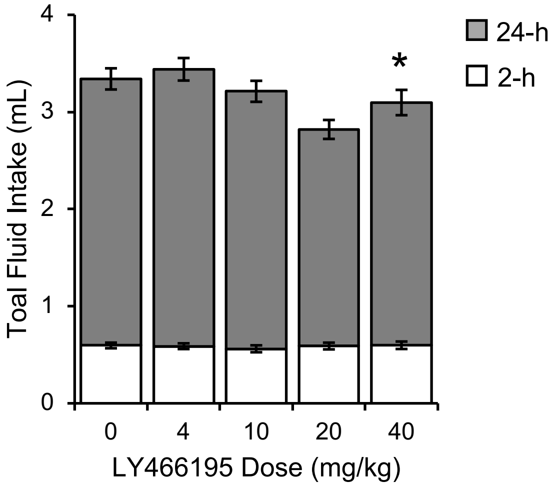

**Figure S2.** Moderate doses of LY466195 do not affect voluntary sucrose consumption

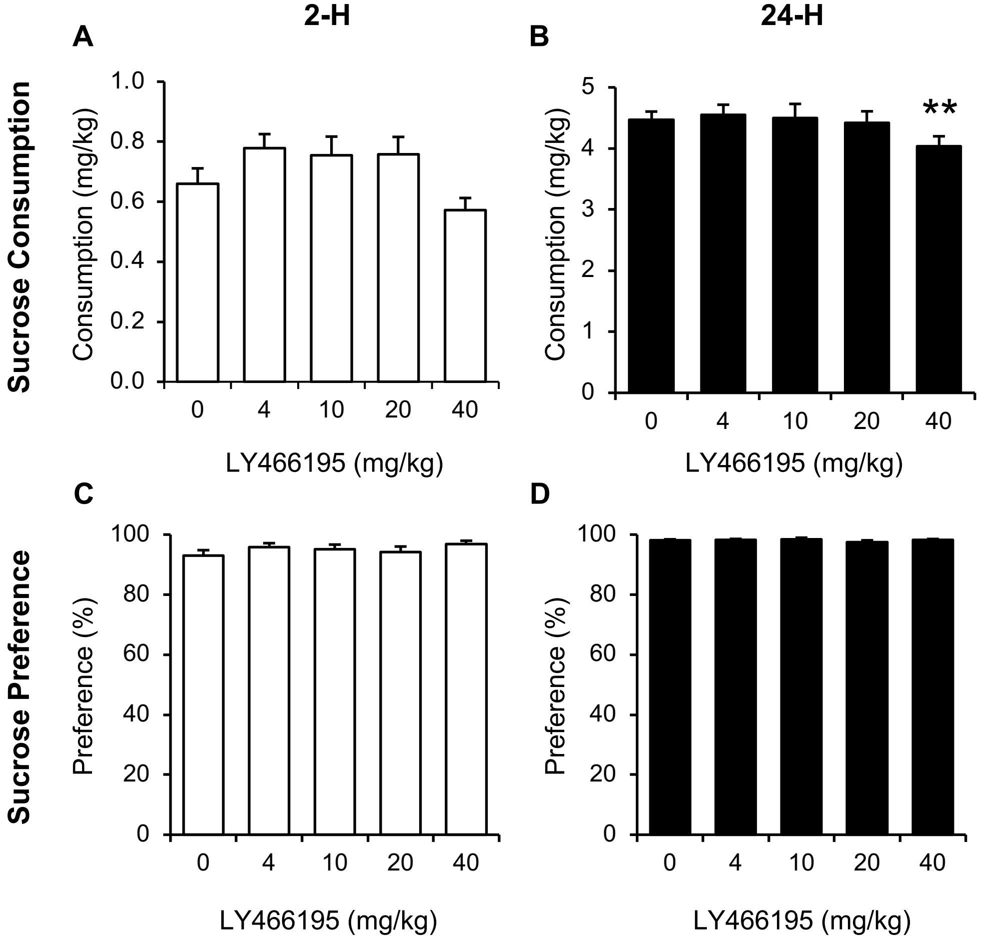

**Figure S3.** Acute LY466195 promotes a transient, dose-dependent effect on voluntary saccharin consumption

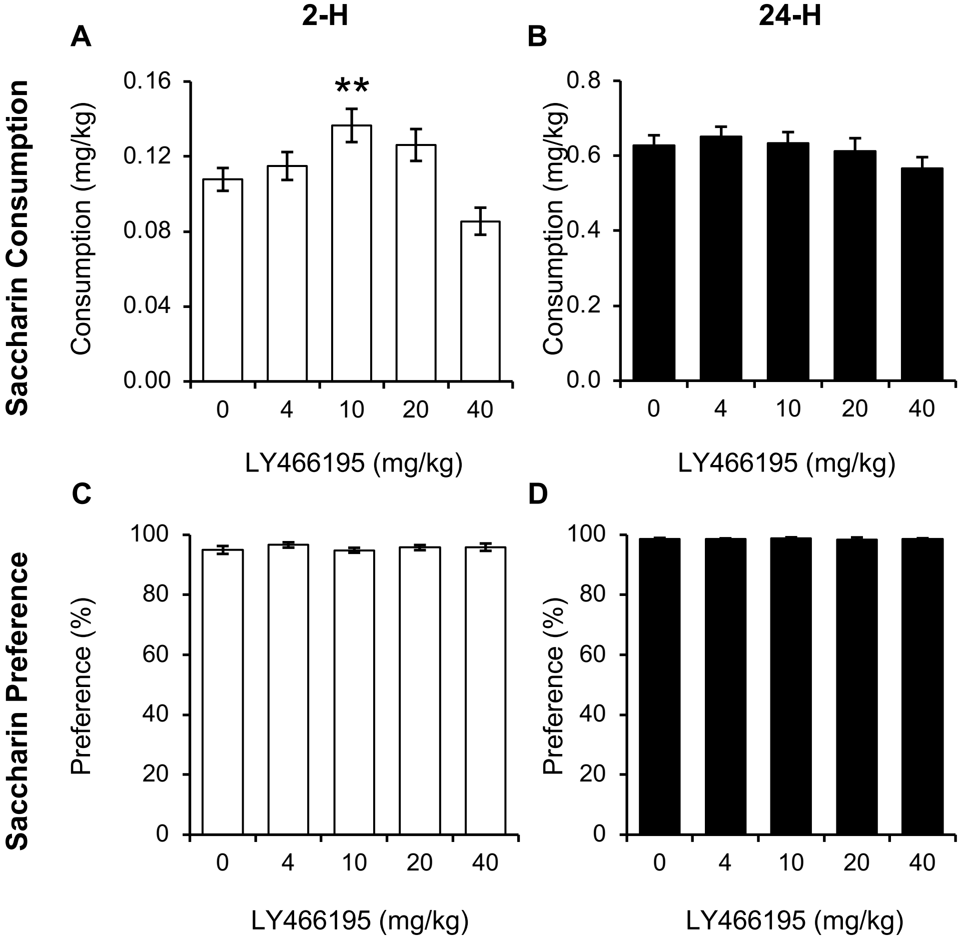

**Figure S4.** Acute LY466195 does not affect voluntary quinine consumption

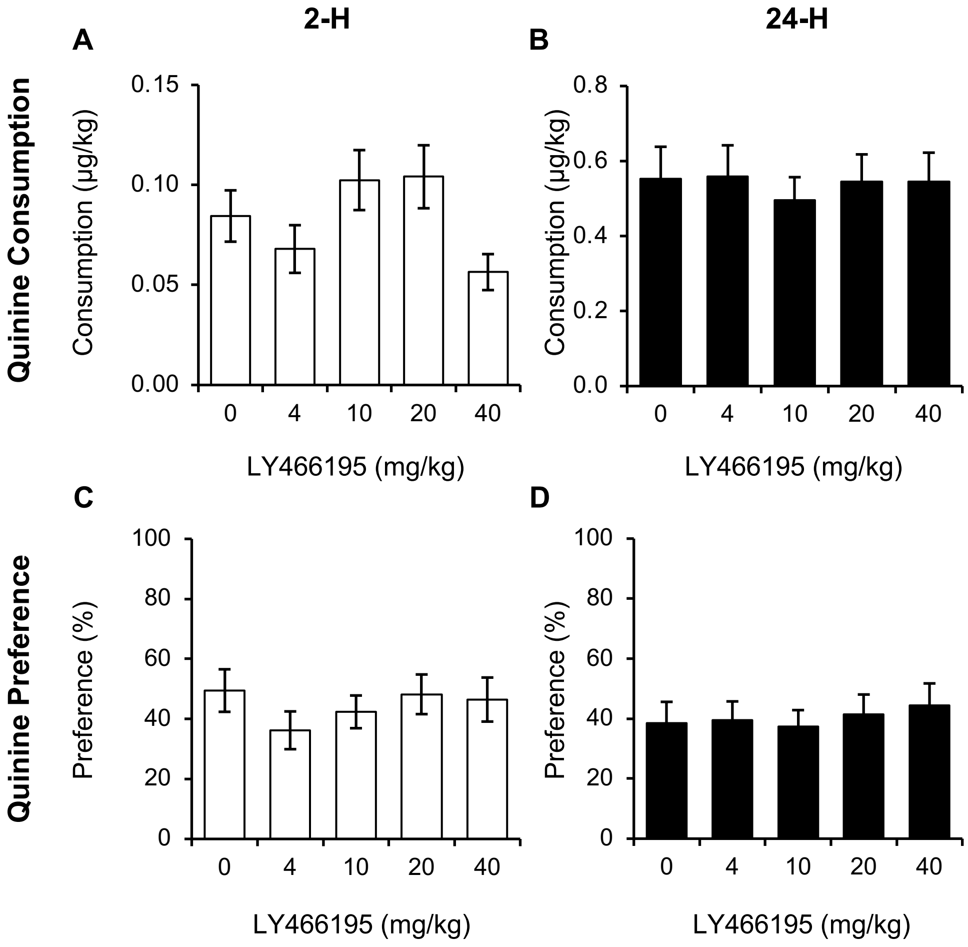

**Figure S5.** Ethanol-related accumbal DA overflow is similar in naïve and ethanol-satiated mice after selective GluK1*KAR blockade

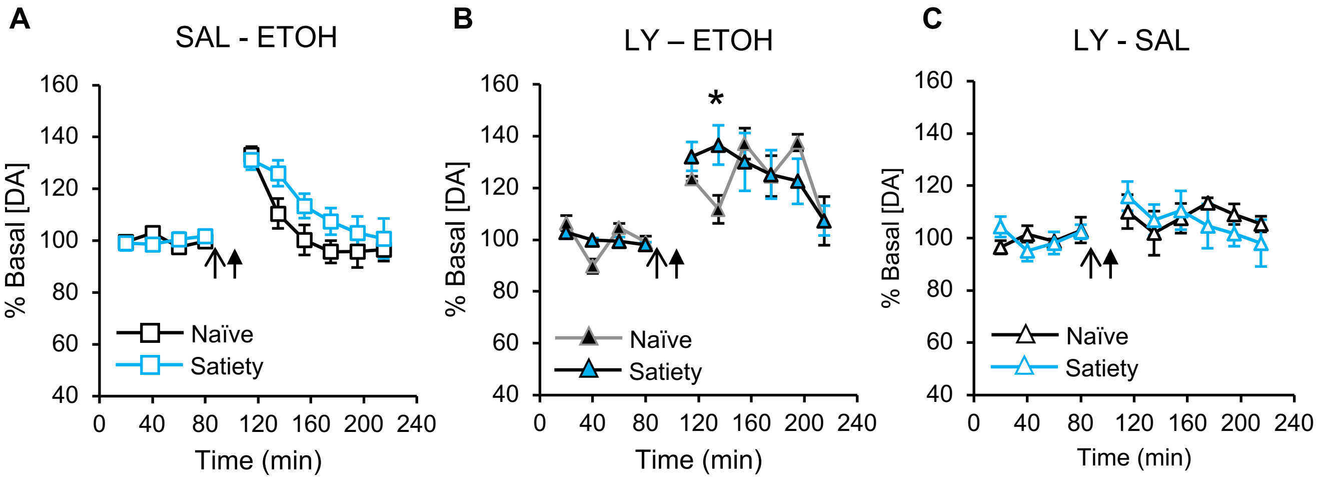

**Figure S6.** Acute LY466195 administration increases locomotor activity in the OFA

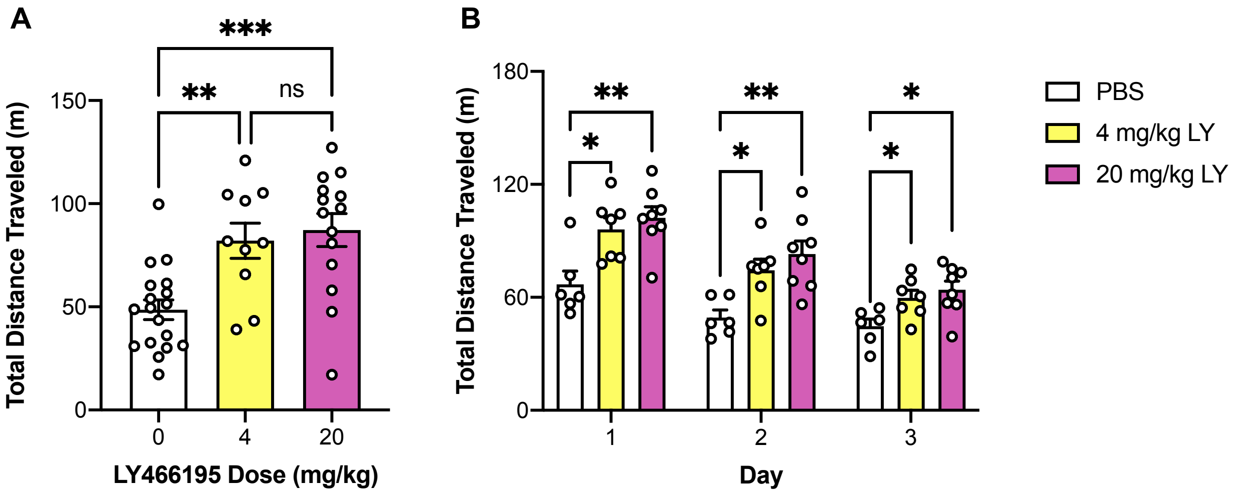
